# Prefrontal cortical ChAT-VIP interneurons provide local excitation by cholinergic synaptic transmission and control attention

**DOI:** 10.1101/461723

**Authors:** Joshua Obermayer, Antonio Luchicchi, Sybren F. de Kloet, Huub Terra, Bastiaan Bruinsma, Tim S. Heistek, Oissame Mnie-Filali, Christian Kortleven, Tim Kroon, Allert J. Jonker, Ayoub J. Khalil, Roel de Haan, Natalia A. Goriounova, Wilma D.J. van den Berg, Christiaan P.J. de Kock, Tommy Pattij, Huibert D. Mansvelder

**Affiliations:** Department of Integrative Neurophysiology, Center for Neurogenomics and Cognitive Research (CNCR), Vrije Universiteit, Amsterdam Neuroscience, The Netherlands; Department of Anatomy and Neurosciences, section Clinical Neuroanatomy, UMC Amsterdam, The Netherlands

**Keywords:** Cortical interneurons, VIP, Acetylcholine, Sustained attention, prefrontal, ChAT, Basal forebrain

## Abstract

Neocortical choline acetyltransferase (ChAT)-expressing interneurons are a subclass of vasoactive intestinal peptide (ChAT-VIP) neurons of which circuit and behavioural function are unknown. It has also not been addressed whether these neurons release both neurotransmitters acetylcholine (ACh) and GABA. Here, we find that in the medial prefrontal cortex (mPFC), ChAT-VIP neurons directly excite interneurons in layers (L)1-3 as well as pyramidal neurons in L2/3 and L6 by fast cholinergic transmission. Dual recordings of presynaptic ChAT-VIP neurons and postsynaptic L1 interneurons show fast nicotinic receptor currents strictly time-locked to single presynaptic action potentials. A fraction (10-20%) of postsynaptic neurons that received cholinergic input from ChAT-VIP interneurons also received GABAergic input from these neurons. In contrast to regular VIP interneurons, ChAT-VIP neurons did not disinhibit pyramidal neurons, but instead depolarized fast spiking and low threshold spiking interneurons. Finally, we find that ChAT-VIP neurons control attention behaviour distinctly from basal forebrain ACh inputs to mPFC. Our findings show that ChAT-VIP neurons are a local source of cortical ACh, that directly excite pyramidal and interneurons throughout cortical layers.

## Introduction

The neurotransmitter acetylcholine (ACh) shapes activity of cortical neurons and supports cognitive functions such as learning, memory and attention ^1–3^. Rapid ACh concentration changes in rodent medial prefrontal cortex (mPFC) occur during successful stimulus detection in a sustained attention task ^4,5^. Traditionally, it is assumed that neocortical ACh is released exclusively from terminals of axonal projections whose cell bodies reside in basal forebrain (BF) nuclei ^6,7^. Chemical lesions of cholinergic BF projections impair attention behaviour ^8–12^ and optogenetic activation of BF cholinergic neurons can mimic ACh concentration changes typically observed during attention behaviour ^11^. Nevertheless, neocortical interneurons that express the ACh synthesizing enzyme choline acetyltransferase (ChAT) have been identified over thirty years ago ^13–15^. They form a sparse population with a predominantly bipolar morphology, are more abundantly present in cortical layers 2/3 (L2/3) ^13,14,16^, and express the GABA synthesizing enzyme Glutamate decarboxylase (GAD), vasoactive intestinal peptide (VIP) and calretinin (CR) ^14, 16–19^. These interneurons could form a local source of ACh in the neocortex, but despite molecular, morphological and physiological characterizations, technical limitations thus far prevented a direct demonstration of whether ChAT-VIP interneurons release ACh or GABA or both. Moreover, BF cholinergic neurons that project to the neocortex have been shown to form direct point-to-point synapses with several types of neurons in different layers, thereby modulating their activity on a millisecond time scale ^20–23^. Activation of ChAT-VIP interneurons can slowly alter local activity of glutamatergic inputs to L2/3 pyramidal neurons ^17^, but it is unknown whether ChAT-VIP interneurons do this via direct cholinergic synaptic transmission, or whether they modulate local neuronal activity more diffusely.

Neocortical circuits contain distinct classes of interneurons with characteristic innervation patterns of local cortical neurons ^19, 24–26^. Fast spiking (FS), Parvalbumin-expressing (PV) interneurons perisomatically innervate pyramidal neurons, while low threshold spiking (LTS), Somatostatin-expressing (SST) target more distal regions of dendrites ^27^. GABAergic VIP neurons inhibit PV and SST interneurons, thereby disinhibiting pyramidal neurons ^26,28,29^. Single cell transcriptomic analysis of cortical neurons has shown that distinct subtypes of VIP interneurons exist with unique gene expression profiles ^19,30^. Whether VIP interneuron subtypes are functionally distinct is not known. It is also not known whether ChAT-expressing VIP interneurons show similar innervation patterns, specifically targeting neighbouring PV and SST interneurons, and activating disinhibitory pathways. Here, we address these issues and find that ChAT-VIP interneurons do not form disinhibitory circuits, but directly excite local interneurons and pyramidal neurons in different mPFC layers with fast cholinergic synaptic transmission. In addition, we show that despite their sparseness, activity of ChAT-VIP neurons is required for sustained attentional performance in freely moving animals.

## Results

### L1 interneurons receive fast cholinergic inputs from ChAT-VIP interneurons

Previous studies in mice have shown that activation of ChAT-VIP interneurons increases spontaneous excitatory postsynaptic potentials (EPSPs) in layer 5 pyramidal neurons ^17^. However, it is unresolved whether ChAT-VIP interneurons directly innervate other neurons in the cortex. To address this, we first expressed channelrhodopsin-2 (ChR2) in ChAT-VIP interneurons in the mPFC of ChAT-cre mice (Fig 1A) and recorded from L1 interneurons since these neurons are known to reliably express nicotinic acetylcholine receptors (nAChRs) in other neocortical areas ^31–33^. All brain slice physiology experiments in this study were done in the presence of glutamate receptor blockers (DNQX, 10 μM; AP5, 25μM). We made simultaneous whole-cell patch-clamp recordings of EYFP-positive ChAT-VIP neurons in L2/3 and nearby L1 interneurons in mouse mPFC (Fig 1B). EYFP-positive neurons showed similar morphology, ChAT, VIP, CR, GAD expression patterns (Fig 1A; Supplementary Fig 1) and action potential profiles (Fig 1B) as reported previously ^14,17,18^. Single action potentials in presynaptic ChAT-VIP interneurons triggered by short (1 ms) electrical depolarization of the membrane potential (Fig 1C) induced fast inward currents in postsynaptic L1 interneurons that lasted up to 10 milliseconds. These fast currents were fully blocked by a combination of nicotinic acetylcholine receptor (nAChR) antagonists, DHßE (10 μM), mecamylamine (MEC, 10 μM) and methyllycaconitine (MLA, 100 nM) (Fig 1C, grey trace). Postsynaptic currents occurred time-locked to the presynaptic action potential with an onset delay of about 2 milliseconds (Fig 1C, bottom graph), suggesting synaptic transmission. In the same recordings, we induced action potentials in presynaptic ChAT-VIP neurons by activating ChR2 with one or two brief blue light pulses, which induced similar fast inward currents in the postsynaptic L1 interneurons that were also blocked by nicotinic receptor antagonists (Fig 1D). Overall, we found in three paired recordings of L2/3 ChAT-VIP and L1 interneurons a unitary synaptic connection that was mediated by nAChR currents. Furthermore, in 67% (n=8 of 12) of mouse L1 interneurons fast synaptic inward currents occurred time-locked to ChR2-induced presynaptic APs in ChAT-VIP interneurons (Fig 1D,J).

**Figure 1.**
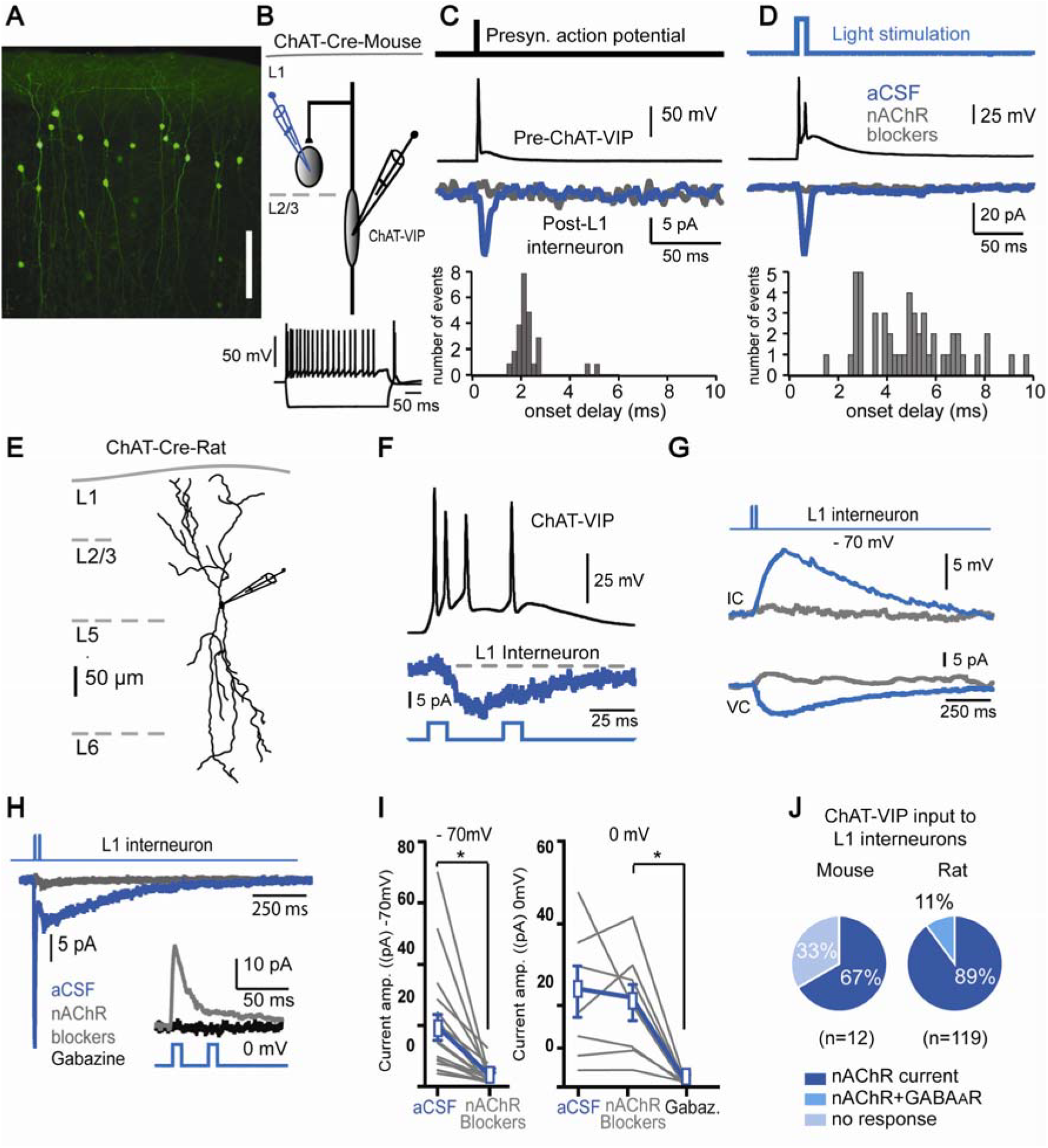
ChAT-VIP interneurons release ACh and GABA. **A)** EYFP-labeled ChAT-VIP interneurons. Labeled cells in L2/3 have predominantly bipolar morphology. **B)** Top: Schematic illustration of the experiment: simultaneous recording of presynaptic ChAT-VIP interneurons and postsynaptic L1 interneurons in mouse mPFC. Bottom: Voltage responses of a L2/3 ChAT-VIP interneuron to depolarizing (+200pA) and hyperpolarizing (−150 pA) somatic current injection. **C)** Example traces of synaptically connected ChAT-VIP and L1 interneuron in the mouse mPFC. Top trace: short step depolarization of ChAT-VIP interneuron to induce an action potential. Middle trace: Presynaptic action potential in ChAT-VIP interneuron. Bottom trace: postsynaptic response of the L1 interneuron showing an inward current (Blue trace) that blocked by nAChR antagonists (DHßE 10 μM, MLA 100 nM and MEC 10 uM, grey trace). Bottom graph: histogram of onset delays of the postsynaptic current relative to the depolarization-induced presynaptic action potential. **D)** Recordings from the same neurons as in **(C)** but now AP firing by the presynaptic ChAT-VIP interneuron was induced by activating ChR2 using a brief blue light flash. Traces and graph as in **(C)**. **E)** Digital reconstruction of an EYFP-positive ChAT-VIP interneuron in the rat mPFC. Scale bar 200μm. **F)** Response to blue light-induced ChR2 activation (470nm, 10ms, 25Hz) of a rat mPFC ChAT-VIP neuron (black trace, top panel, voltage response). Middle trace: postsynaptic response in a simultaneously recorded L1 interneuron. Bottom: blue light stimulation protocol applied. **G)** L1 interneuron is depolarized by blue light ChR2-mediated activation of ChAT-VIP cells. A a single component inward current underlies the depolarization. Both the inward current and the depolarization are blocked by nAChR blockers (MLA 100 nM and MEC 10 μM, grey traces). **H)** L1 interneuron recording showing light-evoked biphasic synaptic input currents at −70mV in aCSF (blue trace) or with nAChR blockers (MLA 100 nM and MEC 10 μM, grey trace). Inset: same L1 interneuron recording at 0 mV showing light-evoked synaptic currents in the presence of nAChR blockers (grey trace) or Gabazine (10 μm, black trace). **I)** Left: current amplitudes at −70mV in aCSF and with nAChR blockers (aCSF: 17.97±4.235 pA, nAChR blockers: 2.55±0.6915 pA, p=0.0012, paired t-test, two-tailed, t=3.888, df=17, n=18). Right: amplitudes recorded at 0mV (aCSF: 20.18±5.794 pA, nAChR blockers: 22.77±7.932 pA, Gabazine: 1.803±0.6177 pA, One-way ANOVA:F_(6, 12)_=2.256, p=0.0220, n=7). **J)** Left: Pie chart showing the percentage of mouse L1 interneurons receiving direct nAChR-mediated synaptic input from ChAT-VIP interneurons. Right: Same as left, but for rat L1 neurons receiving nAChR and GABA_A_R-mediated synaptic inputs from ChAT-VIP interneurons.

In mouse cortex, about 15% of VIP neurons express ChAT ^30^. In contrast, in the PFC of rats about 30% of VIP neurons express ChAT ^18^. To test whether ChAT-VIP neurons more reliably innervate L1 interneurons in rat neocortex, we expressed ChR2 in ChAT-VIP interneurons in the mPFC of ChAT-cre rats ^34^. Following mPFC injections, we did not observe significant retrograde labelling of cells in the basal forebrain (Supplementary Fig 2A,B). In rat prefrontal cortex, EYFP-positive L2/3 ChAT-VIP neurons also had a bipolar morphological appearance (Fig 1E), as reported ^14,18^. Upon activation of ChR2, ChAT-VIP interneurons fired action potentials and simultaneously recorded L1 interneurons showed postsynaptic inward currents (Fig 1F). In all recorded L1 interneurons, blue light activation of ChR2 expressed by ChAT-VIP neurons generated postsynaptic depolarizations and inward currents that were blocked by the mix of nAChR blockers (Fig 1G). These currents were either mono-phasic, consisting of only a fast (Fig 1D) or slow component (Fig 1G), or were biphasic, consisting of a fast and a slow component (Fig 1H), reminiscent of synaptic fast α7-containing nAChR and slow β2-containing nAChR currents expressed by L1 interneurons in sensory cortical areas ^31,32^. The nAChR antagonists MLA and MEC blocked both current components (Fig 1H), showing that in rat mPFC L1 interneurons received direct excitatory fast cholinergic inputs from ChAT-VIP interneurons.

Since ChAT-VIP interneurons can co-express the acetylcholine (ACh) synthesizing enzyme ChAT and the GABA synthesizing enzyme GAD (Supplementary Fig 1) ^17,18^, we asked whether these neurons release GABA in addition to ACh. To test this, the membrane potential of rat mPFC L1 interneurons was held at 0 mV in the presence of nAChR blockers (Fig 1H **inset**). Blue light activation of ChR2-expressing ChAT-VIP cells evoked fast outward currents in 11% of the cells (n=13/119), which were blocked by gabazine (10 μM; Fig 1H,I). In mouse mPFC, we did not observe GABAR currents in layer 1 interneurons (Fig 1J). In rat mPFC, all L1 interneurons received fast cholinergic inputs from ChAT-VIP cells and a minority received both ACh and GABA (Fig 1J).

### No disinhibition of L2/3 pyramidal neurons by ChAT-VIP interneurons

VIP interneurons have been shown to disinhibit L2/3 pyramidal neurons by inhibiting activity of fast spiking (FS), Parvalbumin-expressing (PV) interneurons and low threshold spiking (LTS), Somatostatin-expressing (SST) interneurons ^26,28,29^. To address the question whether ChAT-VIP interneurons form disinhibitory circuits in L2/3, we made whole cell patch-clamp recordings of L2/3 pyramidal neurons and triggered activity in ChR2-expressing ChAT-VIP interneurons by applying blue light pulses. Light-induced activation of ChAT-VIP interneurons did not induce GABAergic synaptic currents in pyramidal neurons, but induced depolarizing inward currents in some pyramidal neurons (n=3 of 18; Fig 2A). These inward currents were blocked by nAChR blockers DHßE, MEC and MLA. Next, we analysed spontaneous inhibitory postsynaptic currents (sIPSCs) received by L2/3 pyramidal neurons. Light-induced activation of ChAT-VIP interneurons did not alter the frequency of sIPSCs received by L2/3 pyramidal neurons (Fig 2B,C), indicating that activity of ChAT-VIP neurons did not change inhibition received by L2/3 pyramidal neurons. To test whether ChAT-VIP target and inhibit other local interneuron types, we recorded from rat mPFC fast spiking (FS, Fig 2D) and low threshold spiking (LTS, Fig 2F) interneurons while triggering activity in ChR2-expressing ChAT-VIP interneurons by applying blue light pulses (Fig 2E,G). We did not observe any GABA-mediated inhibitory postsynaptic currents in both interneuron types following activation of ChAT-VIP interneurons (Fig 2E,G). However, a subgroup of FS (n=4/6) as well as LTS (n=5/8) interneurons showed inward currents at −70 mV that were mediated by fast α7-containing or slow β2-containing nAChRs and were blocked after application of the nAChR antagonists DHßE, MLA and MEC (Fig 2E,G). Thus, we find no evidence that ChAT-VIP neurons form disinhitory circuits in L2/3, as has been reported for other VIP interneurons, but we do find evidence that ChAT-VIP interneurons directly excite subgroups of local interneurons as well as a minority of L2/3 pyramidal neurons.

**Figure 2.**
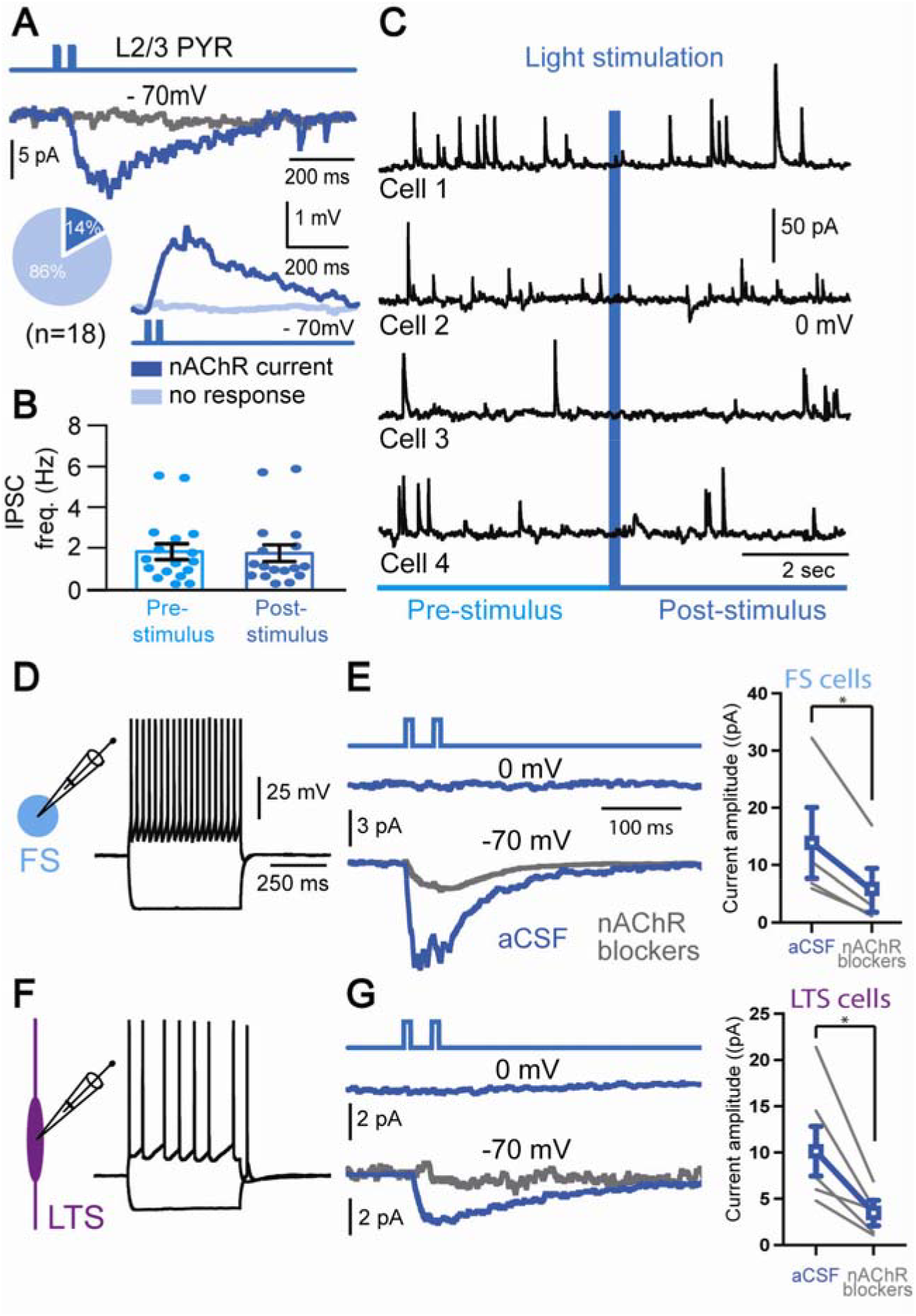
ChAT-VIP interneurons do not disinhibit L2/3 pyramidal neurons. **A)** Example traces of postsynaptic responses in recorded in a L2/3 pyramidal neuron upon ChR2-mediated activation of ChAT-VIP interneurons with blue light (470nm, 10ms, 25Hz, top trace). The postsynaptic responses were blocked by nAChR antagonists (MLA 100 nM and MEC 10 μM, grey trace). The majority of L2/3 pyramidal neurons did not show a postsynaptic response to ChAT-VIP neuron activation (light blue trace). Pie chart showing the percentages of L2/3 pyramidal neurons with nAChR-mediated postsynaptic response (dark blue) and without (light blue). **B)** Comparison of the spontaneous IPSC frequency in L2/3 pyramidal neurons before and after ChR2-mediated activation of ChAT-VIP interneurons with five blue light pulses (25Hz, 10ms),(IPSC frequency pre stimulus: 1.809±0.389Hz, post stimulus: 1.732±0.411Hz, p=0.2310, paired t-test, two-tailed, t=1.245, df=16; n=17; mean±S.E.M.). **C)** Example traces of L2/3 pyramidal neurons recorded at 0mV receiving spontaneous GABAergic IPSCs. Five blue light pulses were applied (25 Hz, 10ms). **D)** Action potential profile of a rat mPFC L2/3 FS interneuron in response to somatic step current injection (+200pA and −150 pA). **E)** Left: Example traces of postsynaptic responses in an FS interneuron upon ChR2-mediated activation of ChAT-VIP interneurons with blue light (470nm, 10ms, 25Hz, top trace). Middle trace: example trace recorded at 0 mV showing absence of an IPSC (n=0/6). Bottom traces: light-evoked postsynaptic currents (n=4/6) in absence (blue trace) or presence of nAChR blockers (MLA 100 nM and MEC 10 μM, grey trace). Right: Summary plot of the postsynaptic current amplitudes of FS cells that showed a response to ChAT-VIP activation. These responses were blocked by nAChR blockers. **F)** Action potential profile of a L2/3 LTS interneuron. **G)** As in **(E)** but for a rat mPFC L2/3 LTS interneuron. No GABAergic IPSCs at 0 mV were observed following light evoked activation of ChAT-VIP interneurons (n=0/8). A subgroup of LTS neurons showed light-evoked postsynaptic currents (n=5/8) at −70 mV (blue trace) that was blocked by nAChR antagonists (MLA 100 nM and MEC 10 μM, grey trace). Right: Summary plot of the postsynaptic current amplitudes of LTS cells that showed a response to ChAT-VIP activation.

### Layer 6 pyramidal receive direct synaptic input from ChAT-VIP interneurons

Previous studies have shown that a majority of layer 6 pyramidal neurons express nAChRs 35,36 and these neurons can be activated by cholinergic inputs from the BF ^21,22,37^. We asked whether L6 pyramidal neurons receive direct inputs from ChAT-VIP interneurons. To test this, we made whole cell patch-clamp recordings from rat mPFC L6 pyramidal neurons combined with activation of ChR2-expressing ChAT-VIP interneurons by applying blue light pulses (Fig 3A,B). Seventy-one percent (n=20/28) of recorded L6 pyramidal neurons showed nAChR antagonist sensitive inward currents (Fig 3B,D). Although the amplitude of these currents was on average about 5 pA, ChAT-VIP activation resulted in a significant depolarization of the membrane potential due to the relatively high membrane resistance of these cells ^22,31^. Six of the L6 pyramidal cells showed an additional gabazine-sensitive fast outward current at 0 mV in the presence of nAChR blockers (Fig 3C,D). These findings show that more than two thirds of the L6 pyramidal neurons receive direct cholinergic inputs from local ChAT-VIP interneurons, and a fifth of L6 pyramidal neurons received both ACh and GABA (Fig 3E).

**Figure 3.**
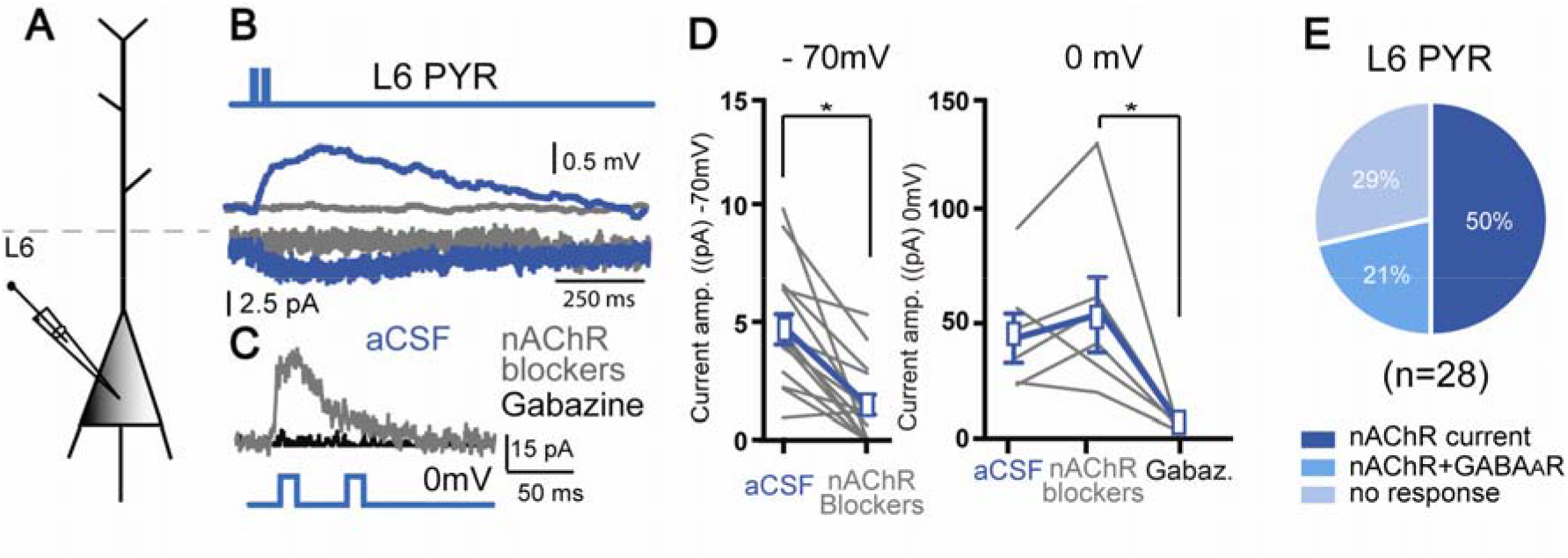
Direct synaptic inputs from ChAT-VIP to L6 pyramidal neurons. **A)** Schematic illustration of recording set up. **B)** Example traces from a rat L6 pyramidal neuron showing depolarization and an inward current at −70mV in response to blue light ChR2-mediated activation of ChAT-VIP neurons (470nm, 10ms, 25Hz) in absence (blue trace) or in the presence of nAChR antagonists (grey trace). **C)** Same L6 pyramidal neuron recorded at 0mV membrane potential showing light-evoked synaptic current in the presence of nAChR blockers (grey trace) and Gabazine (black trace). **D)** Left: summary chart showing the current amplitudes at −70mV membrane potential without and with nAChR blockers (aCSF: 4.820 ± 0.6853 pA, nAChR blockers: 1.483 ± 0.4594 pA, p=0.0002, paired t-test, two-tailed, t=5.051, df=13; n=14, mean±S.E.M.). Right: amplitudes recorded at 0mV with nAChR blockers and Gabazine (aCSF: 40.85 ± 10.35 pA, nAChR blockers: 50.65 ±15.47 pA, Gabazine: 1.403 ± 0.8461 pA, One-way ANOVA:_F(5, 10)_=2.949, p=0.0148; n=6, mean±S.E.M.) **E)** Pie chart showing percentages of L6 pyramidal neurons with nAChR-mediated, combined nAChR and GABA_A_R-mediated, and no synaptic currents.

### Consequences of co-transmission of ACh and GABA

ChAT-VIP cell-induced activation of postsynaptic nAChRs by ACh results in depolarization of postsynaptic cells in L1, L2/3 and L6 as shown above. It is somewhat surprising that some postsynaptic neurons also receive GABA and show inhibitory GABAR currents. Release of GABA in addition to ACh and activation of GABAR currents could lead to shunting inhibition, preventing AP firing. Alternatively, it could result in rebound excitation and augment the excitation provided by nAChR activation ^38,39^. The majority of excitatory nAChR-mediated synaptic responses had slow kinetics with rise times of 155.5±26.5 ms (Fig 4A,C). The subset of combined nAChR and GABAR-mediated postsynaptic responses showed that the GABAergic current had much faster kinetics that decayed back to baseline in about 30 ms (Fig 4B,C). In only four L1 interneurons that showed a combined nAChR and GABAR-mediated current (n=4/13) we found both the fast MLA-sensitive nAChR current that would match the activation kinetics of the fast GABAR current.

**Figure 4.**
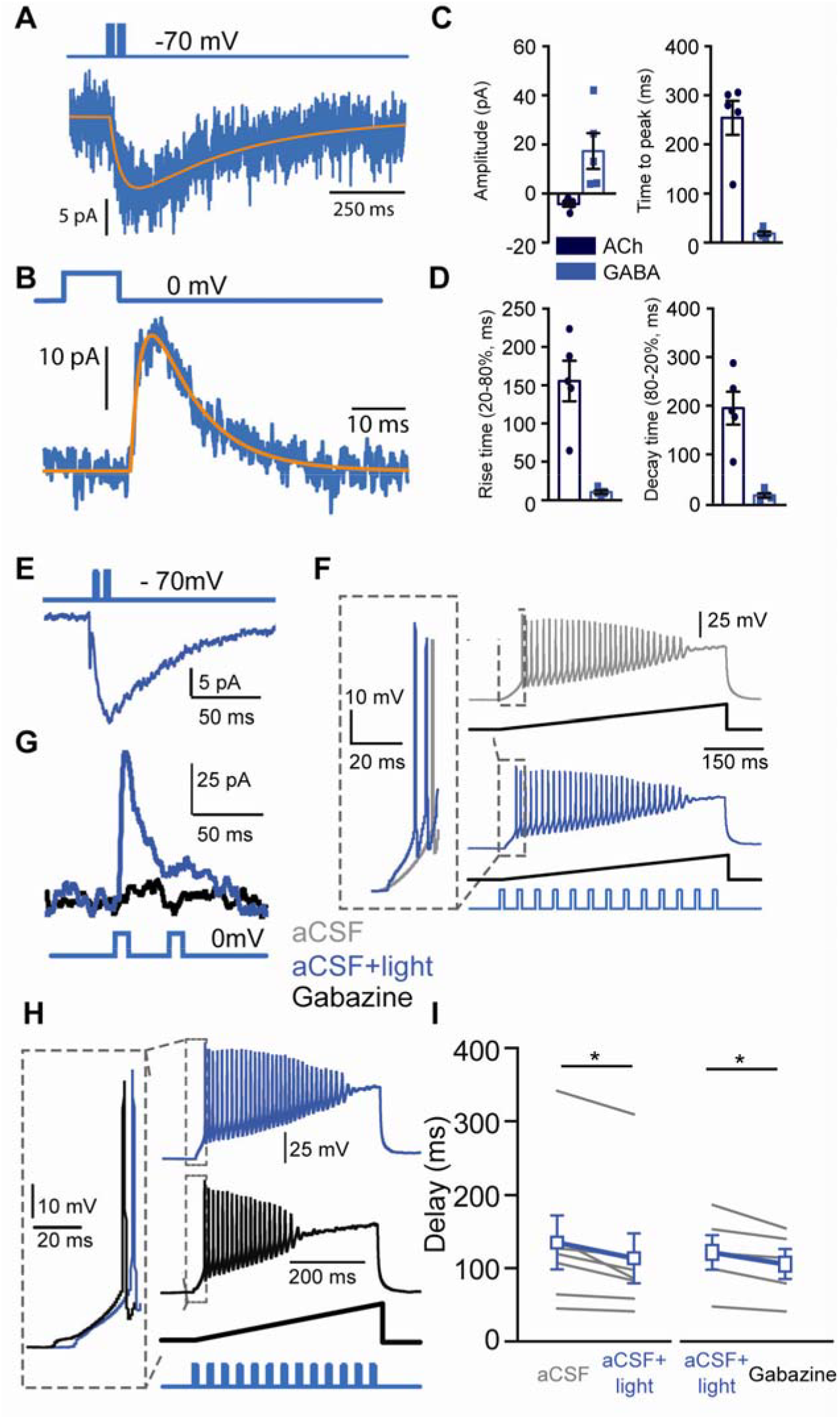
Co-transmission of GABA with ACh postpones AP spiking. **A)** Postsynaptic nAChR-mediated current (blue) recorded at −70mV, with fitted trace (orange). **B)** Same cell at 0mV in the presence of nAChR blockers showing the GABAR-mediated postsynaptic current. **C)** Summary plots of amplitude and time to peak of recorded nAChR and GABAR currents. **D)** Summary plots of rise and decay kinetics of recorded nAChR and GABAR currents. **E)** L1 interneuron as in showing an inward current at −70 mV in response to light-evoked ChR2-mediated activation of ChAT-VIP neurons. **F)** Example traces showing action potential firing in response to a voltage ramp (ramping current injection 1pA/ms for 500ms) in control (grey trace) and with ChR2-mediated activation (13 blue light pulses, 10ms, 25Hz) of ChAT-VIP interneurons (blue trace). **G)** Light-evoked postsynaptic current response in a L1 interneuron held at 0 mV in the absence (blue trace) or presence of Gabazine (black trace). **H)** As in (**F**) but either with blue light stimulation (blue trace) or blue light stimulation in the presence of Gabazine (black trace). **I)** Summary plots of the time to first AP in cells without and with blue light-evoked activation of ChAT-VIP interneurons (Left; aCSF: 136±37.06 ms, aCSF+light: 114.2±34.25, p=0.0159, paired t-test, two-tailed, t=3.326, df=6, n=7) Right: summarizing the time to first AP in cells containing both nAChR and GABA_A_R-mediated postsynaptic currents in absence or presence of Gabazine (aCSF+light: 96.07±18.7 ms, Gabazine: 83.65±16.28, p=0.03, paired t-test, two-tailed, t=3.268, df=4, n=5, mean±S.E.M.).

Hyperpolarizing GABAergic inputs can give rise to rebound excitation by deinactivation of intrinsic voltage-gated conductances ^40^. The excitation induced by slow inward nAChR currents may theoretically be amplified by rebound excitation induced by GABAergic hyperpolarization. To test this, we recorded from L1 interneurons and monitored action potential timing in response to monotonic ramp depolarizations with and without blue light activation of ChR2 expressing ChAT-VIP interneurons (Fig 4E,F). First, to test the effect of the cholinergic component of ChAT-VIP input, only recordings with nAChR-mediated postsynaptic currents without GABAR currents were included (Fig 4E). Activation of ChAT-VIP interneurons advanced the timing of the first AP (Fig 4F), reducing the AP onset delay (Fig 4I). Next, we investigated whether co-transmission of GABA facilitates the advancement of first AP firing, or postpones it. Now, only recordings with combined nAChR/GABAR-mediated postsynaptic currents were included (Fig 4G). Blocking GABAergic inhibition with the GABA_A_ receptor antagonist Gabazine resulted in a shortening of the delay to the first AP in L1 interneurons (Fig 4H), suggesting that the postsynaptic GABAR currents provided shunting inhibition that postponed AP firing. Gabazine did not alter excitability and did not advance spiking in L1 neurons that did not show co-transmission of GABA (**not shown**). In line with these findings, at near-AP threshold membrane potentials in L1 interneurons, blue light activation of ChR2-expressing ChAT-VIP interneurons augmented AP firing probability much more when GABARs were blocked by GABAzine **(Supplemental Fig 3)**. Taken together, these results show that postsynaptic nAChR currents induced by ChAT-VIP interneurons directly excited L1 interneurons, increasing AP firing probability and shortening delays to first AP firing. Co-transmission of GABA provided shunting inhibition, postponing AP firing, rather than facilitating rebound excitation.

**Figure 5.**
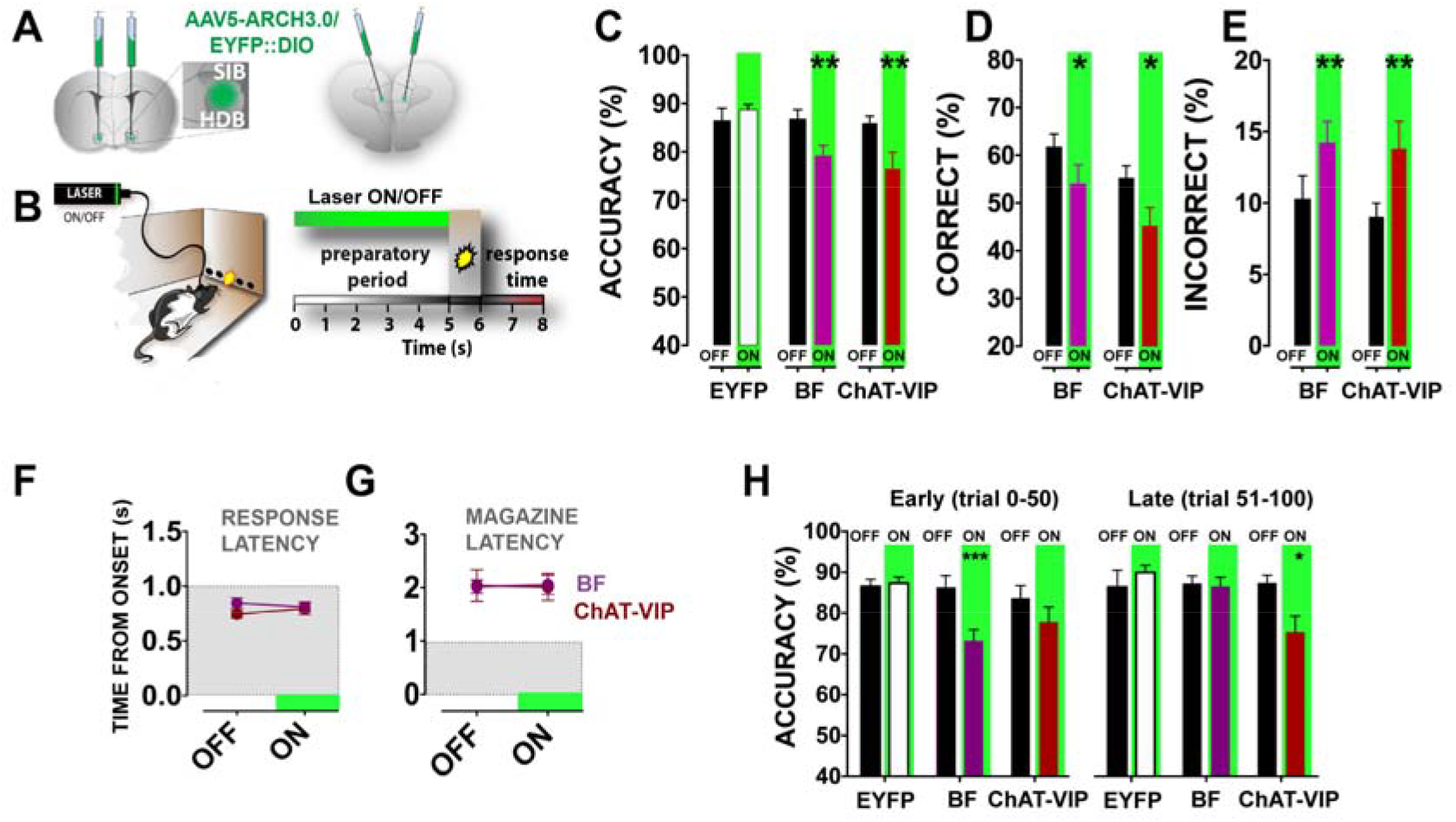
ChAT-VIP interneurons and ChAT-BF neurons control distinct phases of attention. **A)** Locations of virus injections in BF (left) and mPFC (right). In both cases optic fibers were implanted over the PrL mPFC (**Fig S4B**). **B)** Schematic representation of the 5-CSRTT (left) and trails during the task (right) Green bar indicates the period of optogenetic inhibition during the 5 seconds before cue presentation. Laser-on and laser-off trails were assigned randomly in each session (50 ON, 50 OFF). Behavioural performance was constant across multiple sessions on consecutive days (see Methods and Supplemental Figure 4) **C)** Accuracy of responding in rats injected either in BF or mPFC (CHAT-VIP), or control littermates that received only AAV5::DIO-EYFP injections [CTRL: n=9; ChAT-VIP: n=7; BF: n=11; two-way ANOVA, effect of interaction light x virus F_(2,24)_=5.920; p=0.0081; Sidak`s correction ChAT-VIP: p=0.0102 ON vs.OFF; Sidak’s correction BF: p=0.0030 ON vs. OFF, statistics of a single session consisting of 100 trials]. Black bars represent laser-OFF trials. **D)** Percent of correct responses underlying ‘Accuracy’ in **(C)**. (two-way ANOVA; effect of interaction light x virus F_(2,24)_=4.088; p=0.0297; Sidak`s correction ChAT-VIP: p=0.0214, BF: p=0.0288 ON vs. OFF) **E)** Percent of incorrect responses underlying ‘Accuracy’ in **(C)**. (two-way ANOVA; effect of interaction light x virus F_(2,24)_=3.835; p=0.0359; Sidak`s correction ChAT-VIP: p=0.0314, BF: p=0.0254 ON vs. OFF). **F)** Time to respond to light cues and to collect reward at the magazine of the same animals in (**C-E**) [latency correct latency ChAT-VIP: t=1.389; p=0.2141; BF: t=1.576; p=0.142. Incorrect latency ChAT-VIP: t=1.173; p=0.2851; BF: Wilcoxon matched-pairs signed rank test; p=0.6523, ON vs. OFF). Grey shaded area represents the duration of the stimulus light presentation. **G)** As (**F**), ChAT-VIP: Wilcoxon matched-pairs signed rank test; p=0.9063; BF: Wilcoxon matched-pairs signed rank test; p=0.4785; ON vs. OFF). Grey shaded area represents the duration of the stimulus light presentation. **H)** Accuracy of responding during the first half of the session (trails 1-50) and second half of the session (trails 51-100) [two-way ANOVA effect of interaction light x virus: F_(2,24)_=3.744; p=0.0385; Sidak`s correction BF: p=0.0022 ON vs. OFF], ChAT-VIP cells take over its effect in the second half [two-way ANOVA effect of interaction light x virus F_(2,24)_=3.744; p=0.0161 Sidak’s correction ChAT-VIP: p=0.0125 ON vs. OFF]. Green bars represent trails with laser-ON. Black bars represent laser-OFF trials Values are expressed in percent as mean ± S.E.M. *p<.05, **p<.01, ***p<.001.

### ChAT-VIP interneurons are required for attentional performance

Is activity of ChAT-VIP interneurons relevant for mPFC function? To test this, we optogenetically inhibited ChAT-VIP cells during a well-validated task for quantifying attention behaviour, the 5 choice serial reaction time task (5CSRTT) 41 (Fig 5; Supplemental Fig. 4). Since ChAT-VIP cells release ACh in the mPFC, similar to basal forebrain (BF) cholinergic inputs, we also tested whether inhibiting ChAT-VIP interneurons affected attention behaviour distinct from inhibiting BF cholinergic inputs to mPFC. Therefore, ChAT::cre rats received AAV5::DIO-EYFP-ARCH3.0 (or AAV5::DIO-EYFP in controls) injections either in the mPFC or the BF and optic fibers were placed over the PrL mPFC in all groups (Fig 5A; Supplemental Figs 2C-E, and 4). By randomly assigning half of the trials to green laser light ON and the other half to laser OFF (50 trials each), ChAT-VIP cells or BF-to-mPFC projections were either free to fire action potentials or were inhibited in the same animals for five seconds during the pre-cue period when rats show preparatory attention for the upcoming stimulus presentation (Fig 5B). For each animal, behavioural performance during laser ON trials was compared to its own behavioural performance during laser OFF trials. Inhibiting ChAT-VIP cells or BF-to-mPFC projections impaired response accuracy (Fig 5C), and both inhibition of BF cholinergic neurons as well as inhibition of ChAT-VIP interneurons reduced correct responses and increased errors in each animal (Fig 5D,E). Interestingly, no changes in any of the other behavioural parameters were observed, including motor behaviour or motivation to respond as quantified by their response latency and latency to collect the reward (Fig 5F,G; Supplemental Fig 4), These results show that the activity of BF cholinergic projections to the mPFC and the activity of local ChAT-VIP cells are required for proper attention performance.

Interference with the cholinergic system can produce fluctuations in attentive engagement in 5CSRTT ^42,43^. Analysis of attention performance in distinct temporal phases of 5CSRTT sessions showed that inhibiting cholinergic BF-to-mPFC projections reduced accuracy of responding only during the first half of the session (early trials 0-50), but not the second half of the session (late trials 51-100) (Fig 5H). In contrast, inhibiting mPFC ChAT-VIP interneurons significantly reduced attention performance in the second half of the session (Fig 5H). These results indicate that BF ChAT neurons and mPFC ChAT-VIP interneurons affect attention performance distinctly: BF cholinergic neurons support early phases of attention performance, while activity of mPFC ChAT-VIP interneurons is required to sustain attention during the late phase of the session.

## Discussion

In this study, we asked how cortical ChAT-VIP interneurons affect local circuitry in the mPFC, whether they function similar to other cortical VIP cells and whether they are involved in attention behaviour. We found that ChAT-VIP interneurons release ACh locally in both mouse and rat mPFC and directly excite interneurons and pyramidal neurons in different layers via fast synaptic transmission. In contrast to regular VIP interneurons, this ChAT-expressing subtype of VIP interneurons does not inhibit neighbouring fast spiking and low threshold spiking interneurons. Our experiments revealed that activity of ChAT-VIP interneurons contributes to attention behaviour in a distinct manner from activity of basal forebrain ACh inputs to mPFC: ChAT-VIP neurons support sustained attention. These findings challenge the classical view that behaviourally relevant cholinergic modulation of neocortical circuits originates solely from BF cholinergic projections in rodent brain ^44^.

### ChAT-VIP interneurons target local circuitry with fast cholinergic transmission

Various reports over the last thirty years identified neocortical ChAT-expressing VIP interneurons and these were suggested as a local source of ACh in the cortex ^13– 15,17,19^. Simultaneous recordings from cortical ChAT-VIP interneurons and pyramidal neurons showed an AChR-dependent increase of excitatory inputs received by pyramidal neurons following high frequency stimulation of ChAT-VIP interneurons ^17^. However, no evidence was found for direct cholinergic synaptic transmission between ChAT-VIP and other neurons in cortical L2/3 ^45^. We took a different approach from previous studies by recording from neuron populations that have strong nAChR expression ^36^, i.e. L1 interneurons and L6 pyramidal neurons in both mouse and rat mPFC, as well as using ChR2-mediated activation of ChAT-VIP neurons. Our result suggest that ChAT-VIP interneurons form fast cholinergic synapses onto local neurons, since in unitary synaptic recordings the delay between presynaptic action potential and postsynaptic response was about 2 milliseconds, suggesting mono-synaptic connections. Therefore, it is unlikely that ChAT-VIP interneurons triggered poly-synaptic events, exciting terminals of BF neurons and triggering ACh release from these terminals. Fast cholinergic inputs from ChAT-VIP neurons are more abundant in rat mPFC L1 interneurons than in mouse mPFC, in line with a larger percentage of VIP cells expressing ChAT in rat cortex ^18^. Von Engelhardt et al. (2007) ^17^ did not observe nAChR currents activated by mouse ChAT-VIP cells in other L3 interneurons, which we did find in rat mPFC. This may be due to species differences or brain region difference in the two studies. Nevertheless, our findings show that in addition to cholinergic fibers from the BF, ChAT-VIP interneurons act as a local source of ACh modulating neuronal activity in mPFC.

### ChAT-VIP interneurons do not form disinhibitory circuits

Regular cortical VIP interneurons disinhibit local pyramidal neurons by selectively inhibiting somatostatin (SST) and parvalbumin (PV)-expressing interneurons ^24,28,29,46^. In contrast, we did not find evidence that ChAT-VIP neurons form disinhibitory circuits. Low-threshold spiking and fast spiking interneurons receive exclusively cholinergic excitatory inputs and no GABAergic inhibitory input from ChAT-VIP interneurons. Prefrontal cortical ChAT-VIP neurons also did not indirectly disinhibit L2/3 pyramidal neurons through excitation of L1 interneurons ^31^.In mouse auditory cortex, fear-induced activation of L1 interneurons by cholinergic inputs from the BF results in feed-forward inhibition of L2/3 FS interneurons and disinhibition of L2/3 pyramidal neurons ^31^. ChAT-VIP interneurons in the mPFC might in principle play a similar role exciting L1 interneurons as BF cholinergic inputs do in mouse auditory cortex. However, in our experiments we did not find evidence that ChR2-mediated activation of ChAT-VIP neurons altered ongoing inhibition and spontaneous inhibitory inputs to L2/3 pyramidal neurons. In contrast, we found that ChAT-VIP interneurons directly targeted a subgroup of L2/3 pyramidal neurons and provided direct excitation to these pyramidal neurons.

Recent anatomical and functional evidence shows that VIP interneurons in rodent brain are morphologically and functionally diverse and that prefrontal cortical VIP cells can directly target pyramidal neurons ^47–49^. Both multipolar and bipolar VIP cells form synapses on apical and basal dendrites of pyramidal neurons in superficial and deep layers and VIP neurons directly inhibit pyramidal neuron firing ^48,49^. Frontal cortical VIP cells rapidly and directly inhibit pyramidal neurons, while they can also indirectly excite these pyramidal neurons via parallel disinhibition. These findings suggest that not all VIP cell subtypes adhere to targeting only other types of interneurons, and regulating cortical activity through disinhibition only. VIP interneurons represent about 15% of all cortical interneurons in mouse brain, and recent RNAseq profiling identified 12 different molecular VIP-positive subtypes, of which 2 types express ChAT ^19,30^. Our findings show that ChAT-VIP interneurons project to both interneurons as well as pyramidal neurons and directly excite them, in contrast to most regular VIP interneurons. Thereby, activation of ChAT-VIP interneurons in L2/3 of the mPFC can lead to increased excitability of inhibitory as well as excitatory neurons.

### Co-transmission of GABA does not facilitate rebound excitation

In mouse brain, cholinergic fibers from BF neurons can co-transmit the excitatory neurotransmitter ACh with the inhibitory neurotranstmitter GABA in the cortex ^38,39,50^. We find here that in rat mPFC, a minority of L1 interneurons (11%) and L6 pyramidal neurons (21%) receive co-transmission of GABA and ACh from ChAT-VIP interneurons. How these two neurotransmitter interact with each other and what the effect on postsynaptic neurons is, was under debate ^39,51^. Nicotinic AChRs show a range of activation kinetics. Heteromeric β2-subunit-containing nAChR currents have relatively slow activation kinetics with 20-80% rise time of 150 milliseconds, while homomeric α7-subunit-containing nAChR currents activate rapidly with time constants of 2.6 milliseconds and decay time constants of 4.9 milliseconds in neocortical L1 interneurons ^20^, comparable to kinetics of synaptic GABAergic currents. This suggests that when the fast α7-subunit-mediated nAChR currents are induced in L1 interneurons by activation of ChAT-VIP cells, the additional GABAR currents that have similar kinetics will shunt the cholinergic depolarization. In case L1 interneurons express only the slower β2-subunit-containing nAChR currents, co-transmission of GABA could augment the excitatory action of ChAT-VIP neurons by rebound excitation. However, depolarizing ramps or near-threshold action potential firing probabilities revealed that GABA acted inhibitory in both cases, decreasing spiking probability. Therefore, co-transmission of GABA in addition to ACh postpones action potential firing in postsynaptic neurons compared to synaptic transmission of only ACh, forcing a temporal window of inhibition followed by excitation.

This scheme of postsynaptic GABAR current and AChR current interaction will depend on the physical mode of release, whether these neurotransmitters are release from the same ChAT-VIP cell and the same terminals or not ^39,50^. In our experiments using wide-field illumination to activate ChR2 on multiple ChAT-VIP neurons simultaneously, we could not distinguish whether ACh and GABA were release from the same nerve terminals or from the same ChAT-VIP neuron even. It is also not known whether GABA and ACh are packaged in the same vesicles or separately. As such, it is not clear whether co-transmission of GABA and ACh occurs from single ChAT-VIP neurons. However, it is unlikely that ChAT-VIP neurons release only GABA, since we never observed isolated postsynaptic responses mediated only by GABARs, whereas eighty to ninety percent of the postsynaptic responses following ChAT-VIP neuron activation consisted of only AChR currents. So, regardless of the mode of co-transmitter release, ChAT-VIP activity results in excitation and increased spiking probability throughout the mPFC layers.

### ChAT-VIP interneurons support sustained attention performance

Cholinergic signalling in the mPFC controls cognitive attention and task-related cue detection ^4,11,45,52^. In contrast to the general view that ACh is solely released in the mPFC from cholinergic projections from neurons located in the BF, we present here evidence that there is a second source of ACh that supports cognitive attentional performance. The different temporal requirements of activity of BF-mPFC projections and ChAT-VIP interneurons in attention suggests that the two sources of cortical ACh interact in shaping cortical network activity during attentional processing. Our findings indicate that activity of cholinergic projections from the BF is required for early phases of attention performance. In contrast, activity of ChAT-VIP interneurons supports later phases of the attention task. Given the sparseness of these neurons, only 15-30% of VIP interneurons express ChAT ^18,30^, it is surprising that inhibition of this small population in a single brain region has an effect on brain function and behaviour. Even though activation of ARCH expressed by ChAT-VIP cells or axons may lead to increased activity ^53^ or suppression of activity, our experiments do show that specific manipulation of these cell populations affect attention.

Recent findings indicate that BF cholinergic neurons are preferentially activated by reward and punishment, rather than attention ^54^. Hangya et al. suggested that the cholinergic basal forebrain may provide the cortex with reinforcement signals for fast cortical activation, preparing the cortex to perform a complex cognitive task in the context of reward. Still, rapid transient changes in ACh levels in the mPFC may support cognitive operations ^55^ and may mediate shifts from a state of monitoring for cues, to generation of a cue-directed response ^11,52^. Since we find that activity of ChAT-VIP neurons is required during sustained attention, it remains to be determined whether ACh release from local ChAT-VIP interneurons is responsible for or contributes to the generation of cue-directed responses.

## Author contribution

HDM, TP, AL and JO designed the study. AL and OMN performed behaviour experiments. AL, JO and OMN performed surgeries, perfusions and anatomy experiments. SDK, HT and BB assisted in the training, behaviour and anatomy experiments. RDH and CDK provided analysis tools and MATLAB scripts. AL, HDM, and TP analyzed the behavioural data. JO, TH, KK, NAG and AJK performed ex vivo electrophysiology experiments. JO and HDM designed and analyzed the electrophysiological data. TK, AJ en WVDB performed immunostaining experiments. JO, AL, HDM and TP wrote the manuscript with input from all other authors.

## Acknowledgments

We thank JN Vaiña MSc for excellent technical assistance, and Dr. K. Deisseroth for the generous sharing of tools and ChAT-cre rats. HDM received funding for this work from the Netherlands Organization for Scientific Research (NWO; VICI grant 865.13.002), ERC StG “BrainSignals,” EU H2020 Framework Programme (Grant Agreement H2020 HBP 720270), EU 7th framework program (EU MSCA-ITN CognitionNet FP7-PEOPLE-2013-ITN 607508). CdK received funding for this work from the Netherlands Organization for Scientific Research (NWO ALW #822.02.013).

## Methods

### Contact for Reagent and Resource sharing

Further information and requests for resources and reagents should be directed to corresponding author HDM at

### Animals

All experimental procedures were in accordance with European and Dutch law and approved by the animal ethical care committees of the VU University and VU University Medical Center, Amsterdam. Mice: experiments were done on acute brain tissue of both female and male ChAT-IRES-Cre mice (JAX laboratory, mouse line B6;129S6-Chattm2(cre)Lowl/J ^56^). Average age at time of injection was 9 weeks; average age at time of sacrifice was 16 weeks. Rats: male ChAT-cre rats (kindly provided by the Deisseroth lab ^34^) were bred in our facility, individually housed on a reversed 12 h light/dark cycle (lights OFF: 7 a.m.) and were 12-13 weeks old at experiment start. Only when assigned to behavioural experiments, rats were food deprived (start one week before operant training, 85-90% of the free-feeding body weight). Water was provided *ad libitum*. In total 59 rats were included in this study.

### Surgical procedures

All coordinates of injection and fiber placements are from the Rat Brain Atlas (Paxinos and Watson) ^57^. Viruses AAV5.EF1a.DIO.hChR2.EYFP; AAV5.EF1a.DIO.EYFP and AAV5.EF1a.DIO.eARCH3.0 (titer 4.3-6.0×10^12^/ml) were purchased from UPENN Vector Core (Pennsylvania, US). Following anaesthesia (isoflurane 2.5%) and stereotaxic frame mounting (Kopf instruments, Tujunga, USA), the scalp skin was retracted and 2 holes were drilled at the level of either the basal forebrain (BF) or the medial prefrontal cortex (mPFC). Stainless steel micro-needles connected to syringes (Hamilton, USA) were inserted to deliver virus. To optimize rat BF injection location, as we previously did for mouse BF 6, four BF coordinates were used: a) AP −1.20 mm; ML 2.0 mm; DV −6.8 and 8.9 (1μl in total) or −7.8 mm (0.5 μl) from skull; b) AP −0.60 mm; ML 2.0 mm; DV −8.4 mm from skull; c) AP 0.00 mm; ML 1.6 mm; −8.7 and −8.4 (1μl in total) or −8.6 mm (0.5 μl) from skull; d) AP +0.84 mm; 0.9 mm; DV −7.9 and −8.3 (1μl in total) or −8.1 mm (0.5 μl) from skull. For behavioural experiments, injection location in BF was used that resulted in highest EYFP expression in BF to mPFC projection fibers (AP 0.00 mm; ML 1.6 mm; DV −8.7 and −8.4 mm from skull). For mPFC injections were done at AP +2.76 mm; ML 1.35 mm; DV –3.86 and −4.06 mm from skull. For the latter group an infusion angle of 10° was employed ^2^. In all cases, for behavioural experiments 1μL virus was injected per hemisphere in two steps of 500nL, at 6 μL/h infusion rate.

Mice were two to three months of age at time of surgery and virus injection. Analgesia was established by subcutaneous injection of Carprofen (5 mg/kg) and Buprenorphine (100 μg/kg) followed by general anesthesia with Isoflurane (1-2 %). AAV5 virus (EF1a.DIO.hChR2.EYFP) was injected in both hemispheres (400 – 500 nL per hemisphere) of the mPFC (coordinates relative to Bregma: AP – 0.4/-0.4; ML - 1.8 mm; DV – 2.4/-2.7) with a Nanoject (Drummond). Mice were sacrificed for experiments at least three weeks post-surgery.

Following virus delivery in rat brain for behavioural experiments, 2 guide screws and 2 chronic implantable glass fibers (200 μm diameter, 0.20 numerical aperture, ThorLabs, Newton, NJ, USA) mounted in a sleeve (1.25 mm diameter; ThorLabs, Newton, NJ, USA) were placed over the Prelimbic mPFC (200-300 μm on average) under a 10° angle ^58^. Finally, a double component dental cement (Pulpdent©, Watertown, USA) mixed with black carbon powder (Sigma Aldrich, USA) was used to secure optic fibers. All surgical manipulations were performed prior to behavioural training and testing.

### Acute brain slice experiments

Coronal slices of rat or mouse mPFC injected with ARCH3.0 or ChR2 were prepared for electrophysiological recordings. Rats (3-5 months old) were anesthetized (5% isoflurane, i.p. injection of 0.1ml/g Pentobarbital) and perfused with 35 ml ice-cold N-Methyl-D-glucamin solution containing (in mM): NMDG 93, KCl 2.5, NaH2PO4 1.2, NaHCO3 30, HEPES 20, Glucose 25, NAC 12, Sodium ascorbate 5, Sodium pyruvate 3, MgSO410, CaCl2 0.5, at pH 7.4 adjusted with 10M HCl. Following decapitation, the brain was carefully removed from the skull and incubated for 10 min in ice-cold NMDG solution. Medial PFC brain slices (350 μm thickness) were cut in ice-cold NMDG solution and subsequently incubated for three minutes in 34°C NMDG solution. Before recordings, slices were incubated at room temperature for at least one hour in an incubation chamber filled with oxygenated holding solution containing (in mM): NaCl 92, KCl 2.5, NaH2PO4 1.2, NaHCO3 30, HEPES 20, Glucose 25, NAC 1, Sodium ascorbate 5, Sodium pyruvate 3, MgSO4 0.5, CaCl2 1M. Standard equipment for whole-cell recordings were used, as previously described ^60^: Borosilicate glass patch-pipettes (3-6 MΩ), Multiclamp 700/B amplifiers (Molecular Devices), and data was collected at 10 kHz sampling and low-pass filtering at 3 kHz (Axon Digidata 1440A and pClamp 10 software; Molecular Devices).

Recordings from animals injected with ChR2 were made at 32°C in oxygenated aCSF containing in mM: NaCl 125, KCl 3, NaH2PO4 1.25, MgSO4 1, CaCl2 2, NaHCO3 26, Glucose 10. In all of these recordings antagonists to block AMPA receptors 6,7-dinitroquinoxaline-2,3-dione (DNQX, 10 μM), receptors (2R)-amino-5-phosphonovaleric acid; (2R)-amino-5-phosphonopentanoate (AP5, 25μM) and muscarinic receptors Atropine (400 nM) were bath applied. For blocking nAChRs the following antagonists were bath applied: Mecamylamine (MEC, 10μM), DHßE (10μM), and Methyllycaconitine (MLA, 100nM). GABAA receptor mediated responses were blocked by bath application of the antagonist Gabazine (10μM). For whole-cell recordings of EYFP-positive ChAT-VIP interneurons and other L2/3 interneurons a potassium-based internal solution was used containing (in mM): K-gluconate 135, NaCl 4, Hepes 10, Mg-ATP 2, K2Phos 10, GTP 0.3, EGTA 0.2. During recordings, ChAT-VIP interneurons were kept at a membrane potential of −70 mV. Whole-cell recordings of L1 interneurons and pyramidal neurons were made using a cesium gluconate-based intracellular solution containing in mM: Cs gluconate 120, CsCl 10, NaCl 8, MgATP 2, Phosphocreatine 10, GTP 0.3, EGTA 0.2, HEPES 10. Interneurons and pyramidal neurons were identified by their morphology under IR-DIC, the distance of the soma to the pia and their spiking profile. Membrane potentials were kept at −70 or 0 mV to investigate nAChR or GABAR currents.

Opsins were activated by green (530 nm, eARCH3.0) or blue light (470 nm, ChR2). Light pulses with the specific wavelengths were applied to the slices by using a Fluorescence lamp (X-Cite Series 120q, Lumen Dynamics) or a DC4100 4-channel LED-driver (Thorlabs, Newton, NJ) as light source. During recordings from brain slices from animals injected with eARCH3.0, 20 sweeps, each 10s apart were applied. One sweep consists of a 1-s long light pulse. The intensity of the light source was adjusted to 1.7, 3, 7 and 17 watts.

### Immunohistochemistry

Brains from AAV5.EF1a.DIO.EYFP-injected ChAT-cre rats were sectioned in 30 μm-thick slices. BF and mPFC slices were stored in PBS overnight and subsequently incubated in citrate buffer pH 6.0 for 3x 10 min. Thereafter sections were incubated with heated citrated buffer with 0.05% Tween-20 at 90°C for 15 min, left to cool down, and subsequently, rinsed with 0.05M TBS. Next, sections were incubated overnight in 0.05 M TBS with 0.5% triton (Tx) containing all 5 primary antibodies as a cocktail at room temperature. After rinsing slices with TBS (3x 10 min), sections were incubated for 2 hours with secondary antibodies in TBS-Tx. Finally, slices were rinsed in Tris-HCL and mounted on glass slides in 0.2% gelatin, dried, mounted with Mowiol (hecht assistant 1.5H coverslips). As controls, single stained adjaced sectioned were included for all 5 labels.

ChAT staining (Supplemental Figure 1) was performed with anti-ChAT raised in goat (1:300, AB144P, Chemicon Millipore, France) and Alexa Fluor-568-conjugated donkey anti-goat (1:400; A11057, Molecular Probe, Fisher Termo Scientific, Waltham, MA). GAD67 staining was performed with primary antibody anti-GAD67 (1:1200, MAB5406 clone 1G10.2, Chemicon Millipore) and visualized using donkey anti mouse alexa 546 (1:400, A10036, Molecular probe). VIP staining was performed with rabbit anti-VIP (1:1200, 20077 ImmunoStar, Hudson, WI) and donkey alexa-anti-rabbit 594 (1:400, A21207 Molecular probe) as secondary antibody. Further, guinea pig-anti-calretinin (1:4000, 214104, Synaptic systems, Goettingen, Germany) together with donkey-anti-guinea pig alexa 647 (1:400, Jackson 706-605-148).

### Cell counts in basal forebrain

To quantify potential retrograde labeling by AAV5 from the mPFC to the BF (Supplemental Figure 2), rats were injected with AAV5-DIO::eYFP either in the mPFC or the BF at the coordinates used for behavioural and physiological experiments. 50 μm slices of the brains were cut using a vibratome (Leica, 1200T, Germany). Slices were stained for eYFP and mounted on glass slides covered by 2% Mowiol, anti-fading mounting agent and cover slip. Images were acquired using a confocal laser scanning microscope (CLSM; Zeiss LSM 510 Meta) with an excitation wavelength of 514 nm (bandpass 530-600 nm). Cell counting was performed using the cell count function of ImageJ.

### Attention behaviour

After one week of recovery from surgery and 1 week of habituation in the reversed light/dark cycle, rats started training in the 5-CSRTT in operant cages (Med Associates Inc., St. Albans, VT, USA). Training and optical inhibition procedures were analogous to our previously published work with minor adaptations ^58^ (Supplemental Fig 4). In short, following the initial training phase, progression was based on individual performance of each rat, and was reduced from 16 s to 1 s. Criteria to move to a shortened stimulus duration were the percentage of accuracy (>80%) and omitted trials (<20%). When the criterion of 1 s stimulus duration was reached animals were moved to the pretesting phase. In the pretesting phase, a green custom-made LED replaced the normal house-light of the operant cages, (<1 mW intensity) to mask reflections by the laser light used for the experiments.

After three consecutive sessions during which rats performed according to criteria with the LED on in the operant cage, three additional baseline sessions were conducted. During these sessions rats were connected to the patch-cable (Doric Lenses, Quebec city, Canada) used to deliver the light into the brain. In this condition, percentage accuracy was above 80%. However, rats often did not show less than 20% omissions within sessions. This was most likely due to the fact that the animals were connected to the optic fiber patch cable and therefore less free to move in combination with the short time window for the animal to respond (i.e. within two seconds after the cue light went off). Therefore, in line with previous work ^58^, the omission criterion was increased to less than 40% omissions.

Following acquisition of baseline performance, rats were assigned to the testing phase where the task comprised 100 consecutive trials with a random assignment of laser ON or laser OFF trials. For the testing phase, the following parameters were acquired and analyzed through a box-computer interface (Med-PC, USA) and custom-written MATLAB scripts (Mathworks): accuracy on responding to cues (ratio between the number of correct responses per session over the sum between correct and incorrect hits, expressed as percentage); *absolute and percentage of correct, incorrect responses and errors of omission; correct or incorrect response latency; latency to collect reward; number of premature and perseverative responses*. Percent of correct, incorrect and omissions were calculated based on the number of started trials ^59^ to allow a more sensitive evaluation of the parameters.

### Optical inhibition during behaviour

To light-activate the opsins *in vivo*, we used a diode-pumped laser (532 nm, Shanghai Laser & Optics Century Co, China) directly connected to the rat optic glass fiber implant. Light was delivered at 7-8 mW from the fiber tip for experiments carried out with eARCH3.0. These stimulation regimens are able to produce a theoretical irradiance Which ranges between 7.59 and 8.68 mW/mm2 (http://web.stanford.edu/group/dlab/cgi-bin/graph/chart.php). Light was delivered according to scheduled epochs by a stimulator (master 9, AMPI Jerusalem, Israel) connected to the computer interface, which semi-randomly assigned the different trials to laser-OFF or laser-ON conditions (50% of each). In the laser-ON condition, light was delivered during the whole preparatory period (5 s) that precedes stimulus presentation. Optical inhibition sessions were repeated 2 times per rat with a baseline session in between to control for potential carry-over effects.

Moreover, reported data for the majority of rats refer to the first two optical inhibition sessions after establishment of stable baseline performance. Power analysis based on the effect size determined the minimal sample size to detect a statistical significance (7 or more) with a power of β=0.9.

### Histological verification

After behavioural testing, brains were checked for fiber placement and viral expression. For this, rats were anesthetized with isoflurane and a mix of ketamine (200 mg/kg i.p.) and dormitol (100 mg/kg i.p.) and then transcardially perfused (50-100 mL NaCl and 200-400 mL PFA 4%). Brains were removed and maintained in 4% PFA for at least 24 h. After that, brains were sliced with a vibratome (Leica Biosystem, Germany) into 50-100 μm coronal sections and mPFC slices were mounted on glass slides covered by 2% Mowiol, anti-fading agent and cover slip. Images were taken with a CLSM (LSM 510 Meta; Zeiss, Germany) with excitation wavelength of 514 nm bandpass filtered between 530-600 nm, and further analyzed using ImageJ (NIH, USA).

### Quantification and Statisitical Analysis

To evaluate behavioural performance between the ARCH3.0 groups and EYFP control group, two-way ANOVAs for repeated measures were performed. Corrected values for multiple comparison with Sidak’s test were used when the interaction between light and virus was significant. In all cases, the ANOVAs were preceded by the Kolmogorov-Smirnov (KS) test for normal distribution. In cases when the KS p-value was >0.05, factorial analysis was performed on the raw data per parameter. In other cases, raw data were first transformed with square-root or arcsin transformation. Analysis of other parameters were performed with student’s t test, Wilcoxon test and always preceded by KS test to check for normal distribution of the sample. Data were analyzed by MATLAB 2016a (Mathworks), Microsoft Excel (Office) and graphs were plotted by GraphPad Prism. In all cases the significance level was p<0.05.

To statistically evaluate the results between nAChR blockers and aCSF conditions in acute slice experiments, two-tailed paired Student’s t-test was employed. To evaluate differences with GABAR blockers two-way ANOVA for repeated measures was used. To quantify the spike delay time and probability two-tailed paired student’s t-test was used. Significance level was set to p<0.05.

---

**Supplemental Figure 1:**
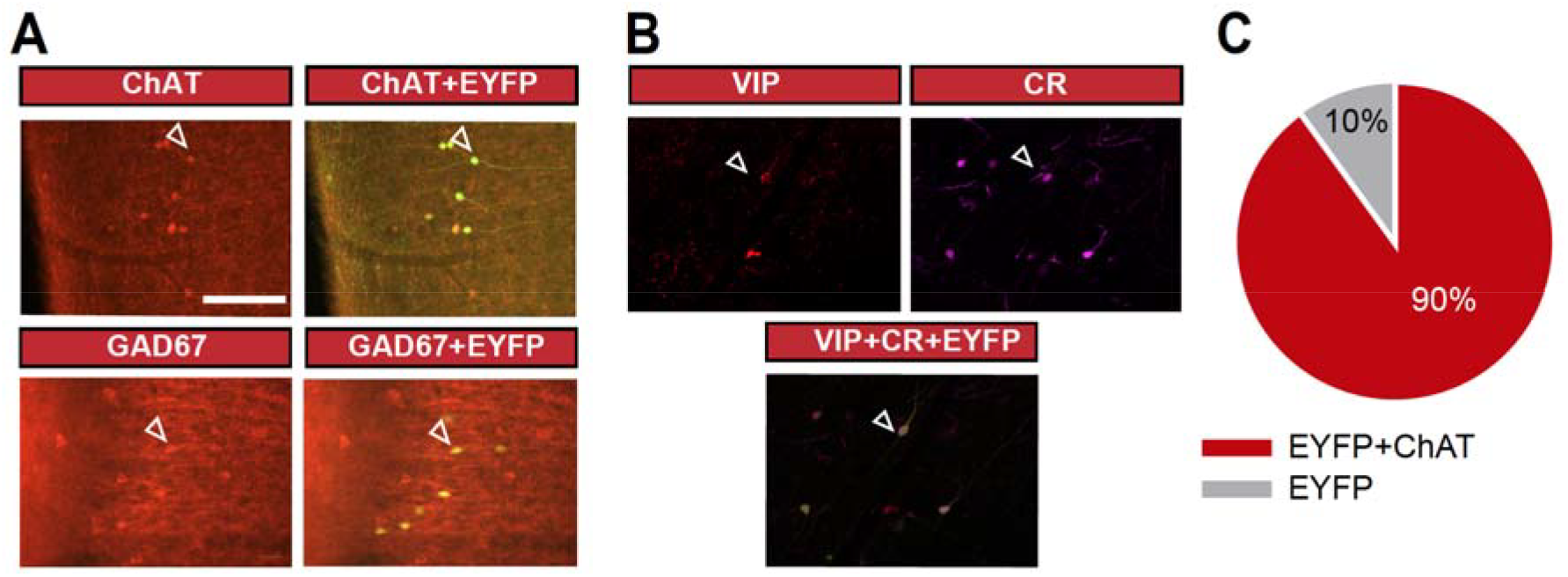
ChAT-VIP interneurons express ChAT, GAD, VIP and CR. **(A)** EYFP-positive interneurons in mPFC of ChAT-cre rats stain positive with antibodies against choline-acetyl transferase ChAT and GABA-synthesizing enzyme GAD67, as reported by Bayraktar et al., 1997 and for mouse by Von Engelhardt et al., 2007 **(B)** EYFP-positive interneurons in mPFC of ChAT-cre rats also stain positive with antibodies against VIP and CR, as reported by Eckenstein and Baughman, 1984, Bayraktar et al., 1997 and for mouse by Von Engelhardt et al., 2007. **(C)** In mPFC of ChAT-cre rats, 90% of EYFP positive cells were found positive for ChAT antibody staining (n=192, 6 animals).

**Supplemental Figure 2:**
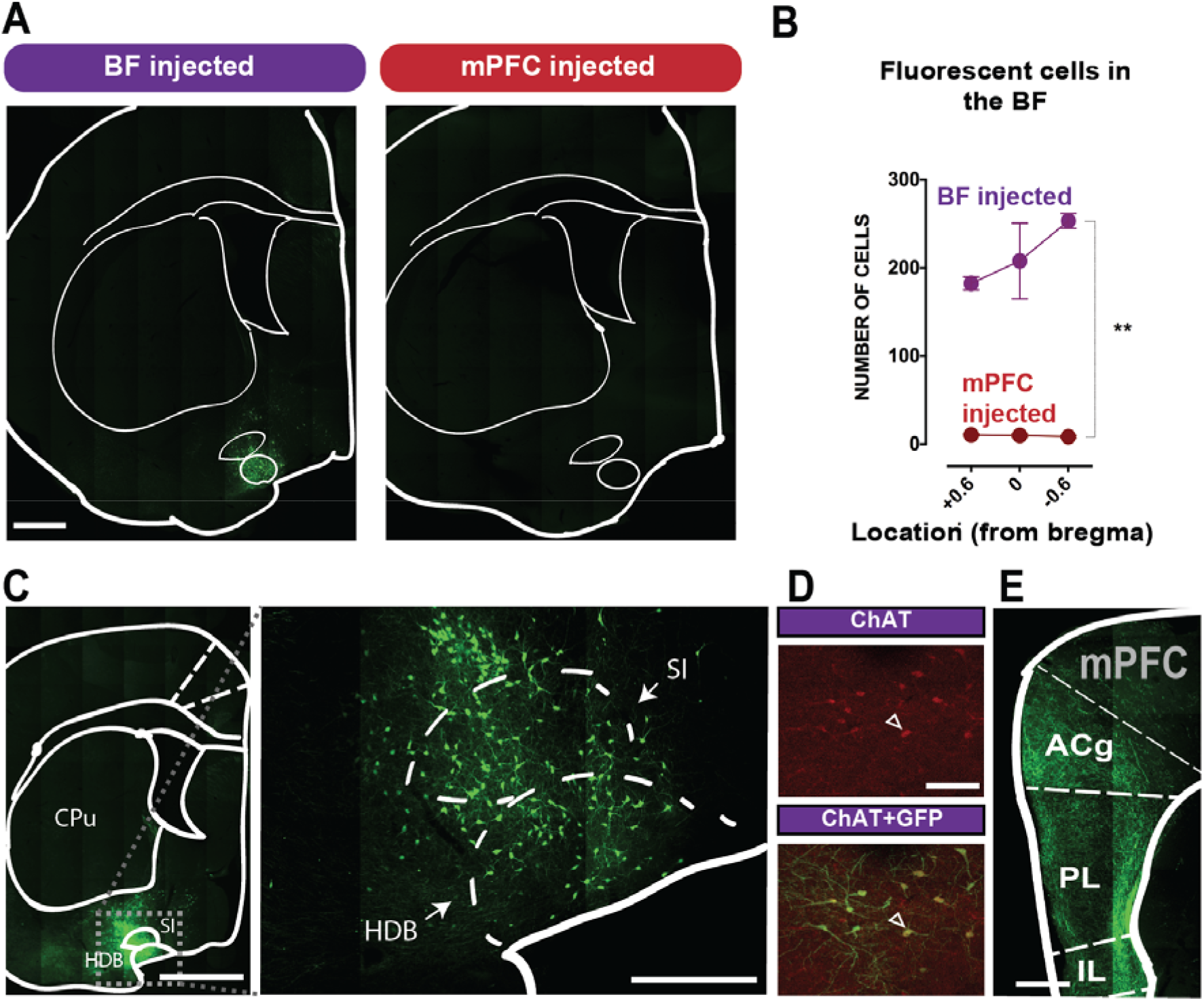
Basal Forebrain cells are not retrogradely labeled by virus injections in the mPFC. **(A)** Basal forebrain (BF) expression of EYFP following AAV5 injection either in BF (left) or in the mPFC (right). **(B)** Number of fluorescent BF cells in BF injected vs mPFC injected ChAT::cre rats [effect of injection location: F(1,2)= 518.1; p=0.001]. Scale bars: 1 mm (1A; 1C left panel); 500 μm (1C right panel); 200μm (E); 70μm (1D). Data are expressed as mean ± S.E.M, ** p<0.01. **(C-E)** AAV5 injections at the level of the HDB and SI (left panel and inset), labels ChAT-positive neurons and fibers in the mPFC. **HDB: horizontal limb of diagonal band of Broca** **SI: substantia innominata**

**Supplemental Figure 3:**
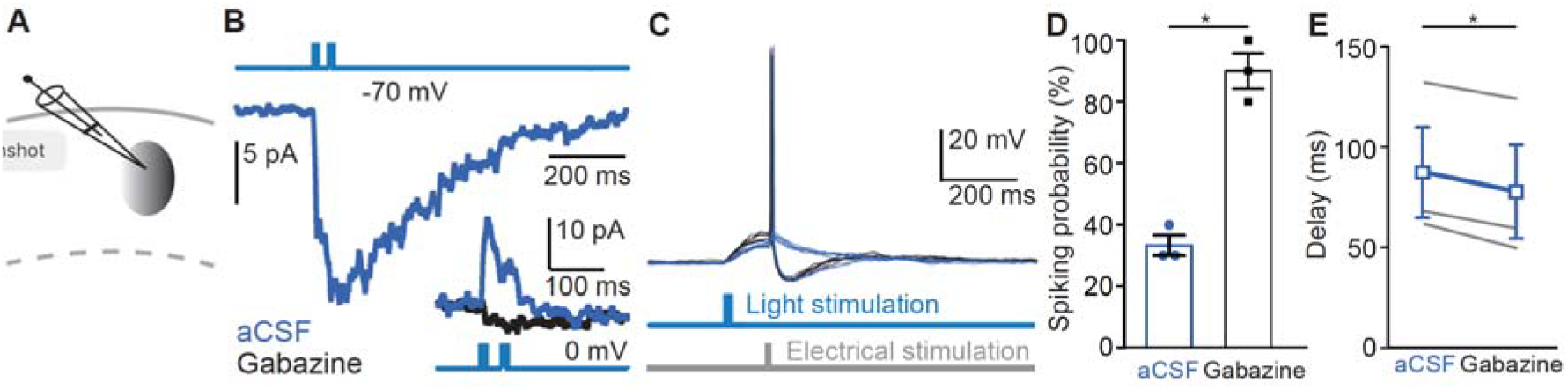
Co-transmission of GABA with ACh reduces spike probability of L1 interneurons. **(A)** Schematic representation of the recording set up. **(B)** Light-evoked ChR2-mediated activation of ChAT-VIP cells generates an inward postsynaptic current response in a L1 interneuron recorded at −70 mV. Inset: same L1 interneuron recording showing a light evoked response at 0 mV in aCSF (blue trace) or Gabazine (black trace). **(C)** Example traces of a recording from a layer 1 interneuron. A short electrical stimulation (1ms) was combined with light evoked activation of ChAT-VIP interneurons. The electrical stimulation was adjusted in that way that the firing probability was ~30% in aCSF (blue trace). The spiking probability was measured again following wash in of Gabazine (black trace) **(D)** Summarizing the spiking probability in aCSF or Gabazine condition (aCSF 33.33±3.33%, Gabazine 90±5.77%, p=0.0234, paired t-test, two-tailed, t=6.425, df=2, n=3). **(E)** As in (**D**) summarizing the time to the first spike (aCSF 87.31 ±22.42ms, Gabazine 77.82 ±23.27ms, p=0.0179,paired t-test two-tailed, t=7370, df=3, n=3, mean ± S.E.M)

**Supplemental Figure 4:**
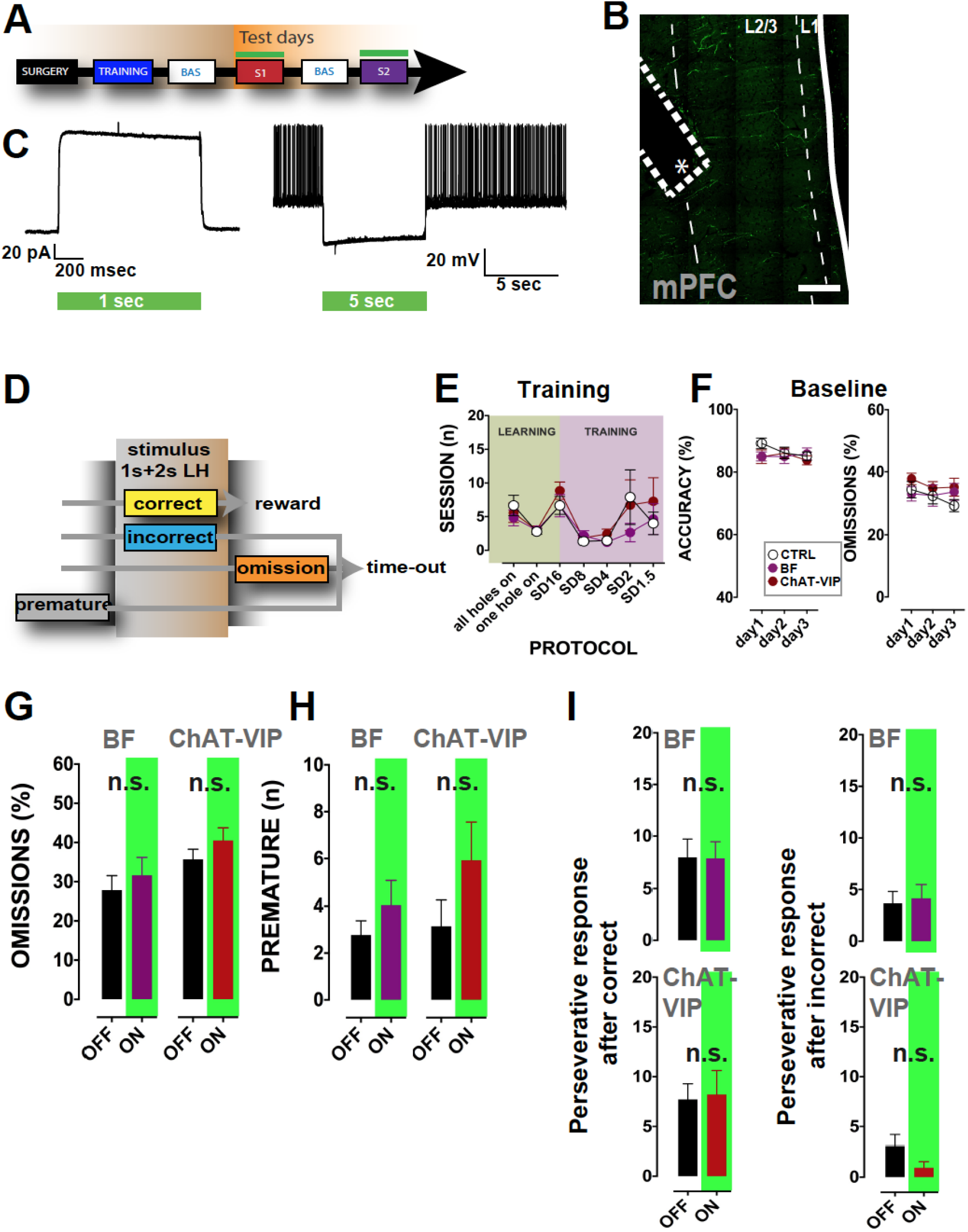
ChAT-VIP or BF projection inhibition during the 5 choice serial reaction time task (5-CSRTT) does not affect omissions, impulsive or compulsive responses. **(A)** Timeline of the behavioral experiments. Following surgery and recovery, rats were trained in the standard version of the 5-CSRTT to stable baseline performance. Next, rats were tested for two sessions with random exposure to green light stimulation in the mPFC (S1 and S2). Between S1 and S2 rats underwent a session of the task without any laser light to test for potential carry-over effects of the light during the former session (BAS). **(B)** Optic fiber location for behavioral experiments. The asterisk indicates the optic fiber tip. In all animals, optic fibers were placed at the border of L2/3 and L5 of the mPFC. Scale bar: 200 μm. **(C)** Whole-cell patch clamp experiments in acute brain slices of ChAT::cre rats injected with AAV5::DIO-ARCH3.0-EYFP used in behavioral experiments show prolonged inhibition upon green light stimulation. Left: voltage-clamp recording of a ChAT-VIP interneuron shows a sustained inhibitory current upon green light exposure. Right: green light evoked hyperpolarization suppressed spiking activity. **(D)** Response types during a trial in the 5-CSRTT. Only correct hits are rewarded with food pellets while all the other responses received 5 second time-out period. **(E)** Neither training duration across the different steps, nor the baseline (accuracy and omission, **F**) differ between the 3 groups (see inset in **F**). **(F)** Neither training duration across the different steps, nor the baseline (accuracy and omission, **F**) differ between the 3 groups (EYFP control, mPFC injected labeling ChAT-VIP neurons, BF injected). **(G)** Errors of omission did not differ when comparing laser-OFF and laser-ON trials, suggesting that both the BF and the ChAT-VIP interneurons play a negligible role in motivational aspects related to attentional performance. **(H)** Similar for premature responses **(I)** and perseverative responses following correct trials **(J)** and perseverative responses following incorrect trials, which were not different in laser-OFF and laser-ON trials.

